# Direct capture and sequencing reveal ultra-short single-stranded DNA in biofluids

**DOI:** 10.1101/2021.08.22.457273

**Authors:** Lauren Y. Cheng, Peng Dai, Lucia R. Wu, David Yu Zhang

## Abstract

Direct capture and sequencing revealed a new DNA population in biofluids, named ultra-short single-stranded DNA (ussDNA), which was neglected by conventional DNA extraction and sequencing approaches. Evaluation of the size distribution and abundance of ussDNA in biofluids manifested generality of its presence in human, animal, and plants. Red blood cells were found to contain abundant ussDNA with enriched functional elements, and thus possesses great potential as a novel DNA biomarker.

## Main

To date, liquid biopsy predominantly focuses on cell-free DNA (cfDNA), which has a prominent peak at 167bp, representing the size of double-stranded DNA (dsDNA) chains wrapped around a nucleosome unit^1, 2^. CfDNA and tissue biopsy testing have established high concordance in advanced cancers^3, 4^; however, cfDNA concentration is low, especially in non-cancer individuals^5, 6^, limiting its application outside of oncology. Moreover, recent studies on cfDNA demonstrate that circulating tumor DNA (ctDNA), the tumor-derived portion of cfDNA, are shorter compared to cfDNA^7, 8^. Another study shows that single-stranded library preparation method has superior cfDNA yield, especially in sub-nucleosomal sizes^1^. These findings suggest that circulating DNA may exist at sizes below nucleosomal size and are likely single-stranded or partially single-stranded. However, conventional DNA extraction methods based on spin column or Solid Phase Reversible Immobilization (SPRI) are susceptible to high loss at sizes below 100nt^9, 10^. Thus, a method tailored for extraction of ultra-short single-stranded DNA (ussDNA) is necessary to uncover the sub-nucleosomal space and explore the presence of biomarkers in various biofluids.

Here, we present a method that characterizes ussDNA from biofluids by direct capture and sequencing, in which probe captured ussDNA are prepared for next generation sequencing (NGS) using single strand-based library preparation (**Fig. 1a**). Probes for direct capture are composed of a pool of oligos with 10-mer degenerate locked nucleic acid (LNA) bases (5’-NNNNNNNNNN-Biotin-3’), and the rationale is that 10-mer degenerate bases bear ∼1 million unique sequences, which can be largely matched to the massively diverse human genome. LNA bases can enhance duplex stability and thus lower melting temperature^11^, so the probe design enables efficient hybridization of diverse populations of ussDNA at room temperature in high salinity buffer condition. The single-stranded library preparation approach leverages a splinter adapter ligation step optimized for recovery of short fragments using organic solvent^12^. Sequencing of 24 different synthetic ssDNA oligos with sizes from 20nt to 70nt showed that all strands were recovered at their expected sizes (**Fig. S1a-b**), albeit 100-fold yield variation was observed following capture (**Fig. S1c-d**), which could be partially explained by highly varied hybridization rates among different sequences^13^. The method demonstrated 9x yield for ssDNA compared to dsDNA as manifested by reads retrieved from equal molarity spike-in (**Fig. S1h**). In addition, an adapter dimer blocker was designed according to a published blocker displacement amplification principle^14^ to reduce dimerized adapters in library (**Fig. S1f-g**).

**Figure 1:**
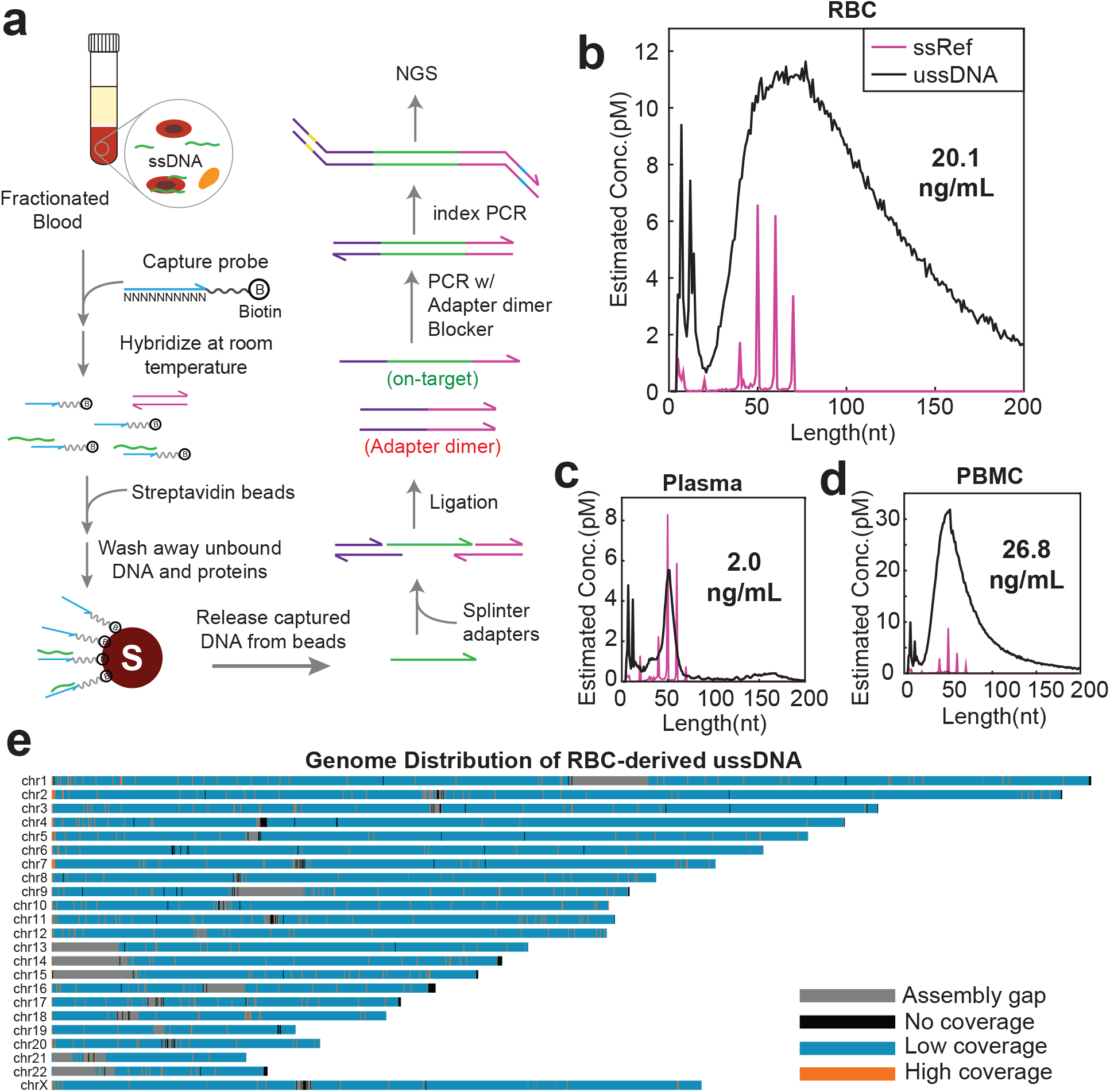
Direct capture and sequencing discovered ussDNA in blood components. **(a)** Direct capture and sequencing workflow. ussDNAs captured by LNA probes were separated from biofluids on streptavidin beads. Extracted ussDNAs were then ligated with adapter sequences and prepared for sequencing. **(b-d)** Length distribution of ussDNA found in **(b)** RBC **(c)** plasma and **(d)** PBMC. Peaks below 20nt that appeared in all samples are artifacts produced by the workflow. Magenta spikes represent ssDNA spike-in references, 4 different strands were spiked in at each size at concentration of 1pM each. **(e)** Human genome distribution of RBC-derived ussDNA in autosomes and chromosome X. Regions where no reads aligned to were shown in black; gaps in human genome assembly GRCh38 were colored in grey. Regions with more than 2x global density were colored in orange.

Next, the direct capture and sequencing approach was applied to plasma, peripheral blood mononuclear cell (PBMC) and red blood cell (RBC) separated from total blood and ussDNAs were found in all blood fractions. RBC-derived ussDNA displayed wide distribution from 20nt to larger than 200nt, with the highest concentration appeared at ∼70nt (**Fig. 1b**); ussDNA from plasma showed a sharp peak at 50nt, and the nucleosome-sized cfDNA was also recovered, likely captured via partial single-strands in dsDNA (**Fig. 1c**); PBMC-derived ussDNA had a similar 50nt peak as plasma ussDNA (**Fig. 1d**). RBC in this healthy individual was found to be abundant in ussDNA, whose concentration is approximately 10-fold higher than plasma, and comparable to PBMC. This finding is counterintuitive because human RBCs are nucleus-free and thus are expected to have low DNA content. Note that when probes were absent from the system, ssDNA spike-in strands were not recuperated in RBC or plasma library, and the 50nt mode in plasma ussDNA did not appear (**Fig. S1i-j**), implying that probes are effective in capturing ssDNA. UssDNA in both RBC and plasma exhibited dominant peaks at 50nt after dsDNase treatment that is specific to dsDNA, and distributions shifted to no-template-control following DNase I digestion, suggesting DNA material with single-strandness at ∼50nt (**Fig. S2a-d**).

To quantitate the concentration of blood ussDNA, ssDNA oligos were spiked into each blood fraction (**Fig. S3a**) at 1pM/strand and the concentration was estimated from the relative abundance of ussDNA and reference sequences. The estimation was validated by curve fitting of varied spike-in concentrations and the corresponding reads fraction of spike-in sequences (**Fig. S3b-c**). In the small cohort of 17 healthy individuals, the approximated ratio of the mean of ussDNA concentration is PBMC:RBC:plasma = 27:7:1 (**Fig. 2a**). Both RBC and PBMC contain significantly more ussDNA than plasma (p<0.0001), despite higher variations in concentration. UssDNA concentrations in different blood fractions had no apparent relationship as RBC ussDNA concentration merely displayed slightly positive correlation with its concentration in plasma and PBMC (**Fig. 2h, Fig. S4f**). Correlation between ussDNA concentration and age was investigated and only the plasma fraction exhibits moderately positive correlation with age (**Fig. 2b-d**). UssDNA concentration in RBC is higher in female than male individuals (p<0.05), whereas no gender difference is manifest in plasma or PBMC counterparts (**Fig. 2e-g**). When spike-in strands were added to total blood prior to fractionation (**Fig. S3d**), the reference strands retained primarily in aqueous phase, i.e., the plasma fraction, and were depleted in cellular phases, especially after washing (**Fig. S3g-k**), inferring that association of bare reference strands with cellular components is not intrinsically favored. In contrast, PBS washing did not deprive RBC of its ussDNA content (**Fig. S3e-f**), suggesting the association of RBC ussDNA with cellular structures.

**Figure 2:**
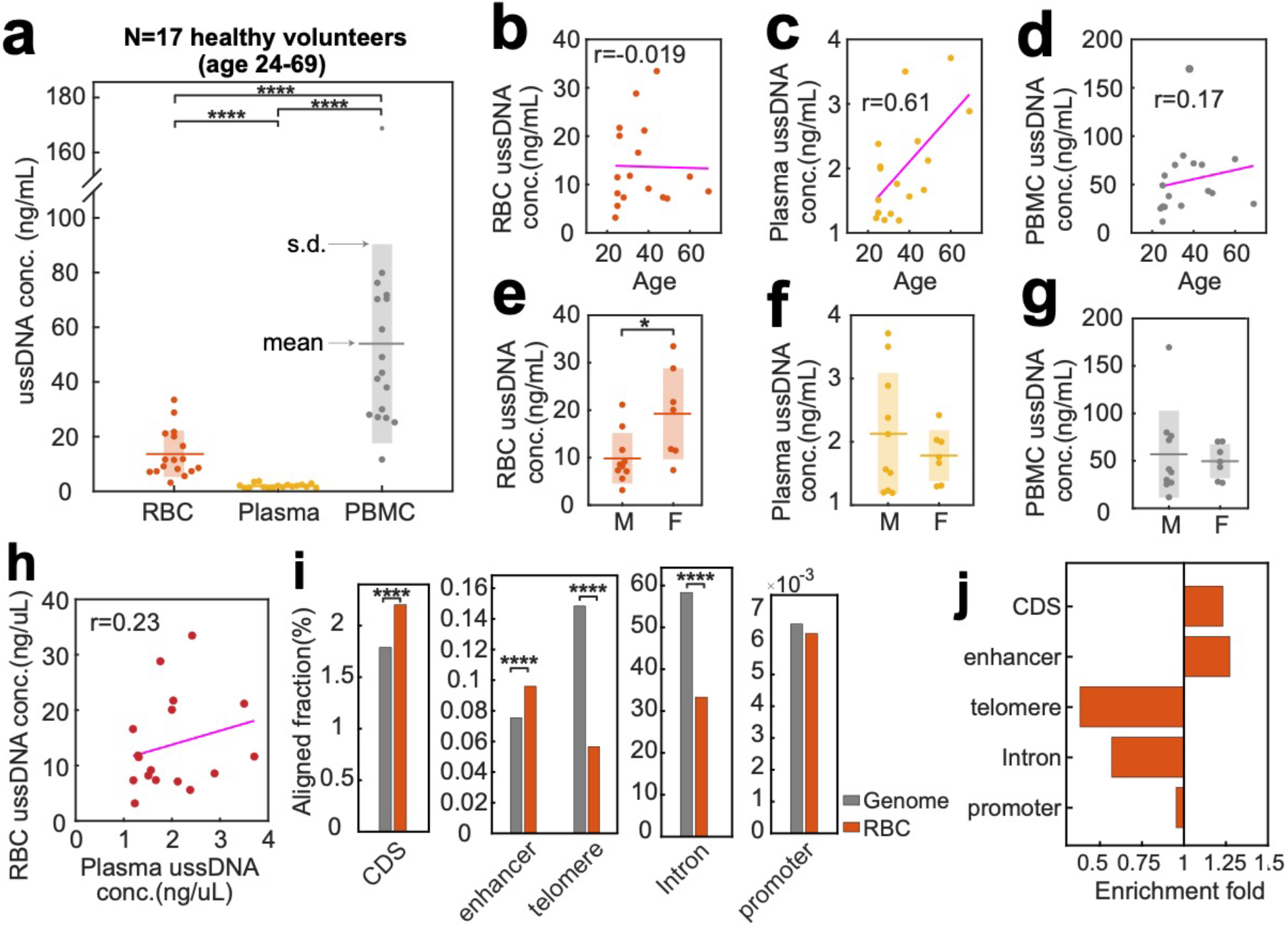
Quantification and focal genome distribution of blood ussDNA. **(a)**ussDNA concentrations in RBC, plasma and PBMC fractions from healthy volunteers. Boxplot represents standard deviation from the mean. ussDNA concentration’s rank is PBMC > RBC > plasma (paired t-test, ^****^ p<0.0001). **(b-d)** Correlation of age and concentrations of ussDNA in **(b)** RBC, **(c)** plasma and **(d)** PBMC. Only plasma ussDNA concentration exhibits moderately positive correlation with age from Pearson’s correlation coefficient r. **(e-g)** ussDNA concentrations grouped by gender (M = male, F = female) in **(e)** RBC, **(f)** plasma and **(g)** PBMC. Welch’s t-test shows significantly higher plasma ussDNA concentration in females compared to males (* p<0.05). **(h)** Correlation between RBC ussDNA concentration and plasma ussDNA concentration. **(i)** Fraction of RBC ussDNA aligned to functional elements compared to their fractions in human genome. CDS: gene coding region. Significant enrichment of enhancer elements and CDS, and depletion in telomere and intron regions are observed in RBC ussDNA compared to random distribution (^****^p<0.0001). **(j)** Fold enrichment of functional elements in ussDNA derived from RBC.

We then sought to understand whether ussDNAs originate from entire or parts of the human genome. Whole genome alignment revealed global distribution in autosomes and chromosome X from RBC-derived ussDNA (**Fig. 1e**). RBC ussDNA distributed uniformly among all chromosomes (**Fig. S4b**). However, 2880-fold enrichment of mitochondria genome was observed in RBC ussDNA, implying exogenous source of ussDNA in RBC fraction because erythrocytes have no mitochondria. The mitochondrial ussDNA in RBC showed increased portion of short fragments compared to nuclear ussDNA (**Fig. S4c**). Although enrichment of mitochondrial cfDNA was reported^6, 15^, the fold enrichment might be partially explained by the fact that human cells contain 200-4000 copies of mitochondria dependent on metabolic intensity^16^. With regards to focal distribution of functional elements, their fractions in reads were compared with corresponding genome fractions. We assumed that the reads fraction is representative of their biological distribution in ussDNA because for highly diverse sequences the biases in hybridization, PCR amplification, or random sampling become negligible. RBC ussDNA showed significant enrichment in gene coding regions and enhancer elements (p<0.0001), as well as depletion in intron and telomere regions (p<0.0001) compared to their distributions in genome (**Fig.2i-j**). The enriched fraction of coding or regulatory regions in RBC ussDNA, in combination with the transporting role of RBCs, may suggest functional roles of RBC ussDNA. Like RBC-derived ussDNA, plasma ussDNA also presented uniformity among all chromosomes despite low coverage owing to lower sequencing depth (**Fig. S4a,d**). Mitochondrial ussDNA in plasma was enriched 430-fold and displayed similar size distribution to its nuclear counterpart (**Fig. S4e**). In plasma ussDNA, depleted intron and telomere regions (p<0.0001) and enriched enhancers (p<0.01) were also observed (**Fig. S4g-h)**.

Last, we expanded our study to biofluids of other animal species and plants to investigate the generality of ussDNA. Unlike RBC ussDNA from human where its distribution is smooth, ussDNA from bovine and pig RBC displayed spiky peaks from 30nt to 70nt with ∼10nt periodicity followed by a smooth decay (**Fig. S5a,d**). UssDNA from bovine plasma showed a similar peak to its human counterpart, except that the peak centered at 63nt instead of 50nt (**Fig. S5b**). Plasmas from pig and rabbit, however, had decreasing oscillations of 10nt periodicity from 30nt to above 100nt (**Fig. S5e,g**). The 10nt periodicity is representative of a helical turn around the nucleosome core, and infers the generation mechanism via DNase I cleavage^1^. Interestingly, the periodic pattern appeared in PBMC of pig (**Fig. S5f**) and milk of bovine (**Fig. S5h**) but not in PBMC of bovine (**Fig. S5c**). To confirm whether animal ussDNA was free of human DNA contamination, bovine ussDNA was cross-aligned to human reference genome and notably reduced coverage was found (**Fig. S6a-d**). Plant-derived ussDNA showed varied sizes and concentrations. In kiwifruit, ussDNA peaked at 34nt followed by gradual reduction (**Fig. S7a**). In orange and cherry, ussDNA spikes between 20-30nt and short artifacts that appeared in all libraries are indistinguishable, and their concentrations above 30nt are low, especially in cherry (**Fig. S7b-c**).

In summary, we present a method tailored to investigate the understudied sub-nucleosomal ussDNA in biofluids. We found ussDNA in blood fractions of human as well as animal species, and plant-based biofluids with different quantities and size distributions. To our surprise, human RBC is rich in ussDNA, which challenges the notion that human RBC do not contain DNA. Enrichment of RBC ussDNA in mitochondrial genome and functional regions such as gene coding region and enhancer elements may imply functional roles of RBC ussDNA. Collectively, our finding may direct to a promising biomarker that has high concentration and won’t subject to interference from nucleus DNA. This study points for the first time to a new population of circulating DNA, yet little is known other than its size representation and concentration, and thus future work is awaiting to elucidate its generation mechanism, tissue-of-origin, disease indications, etc.

## Figures

**Figure S1:**
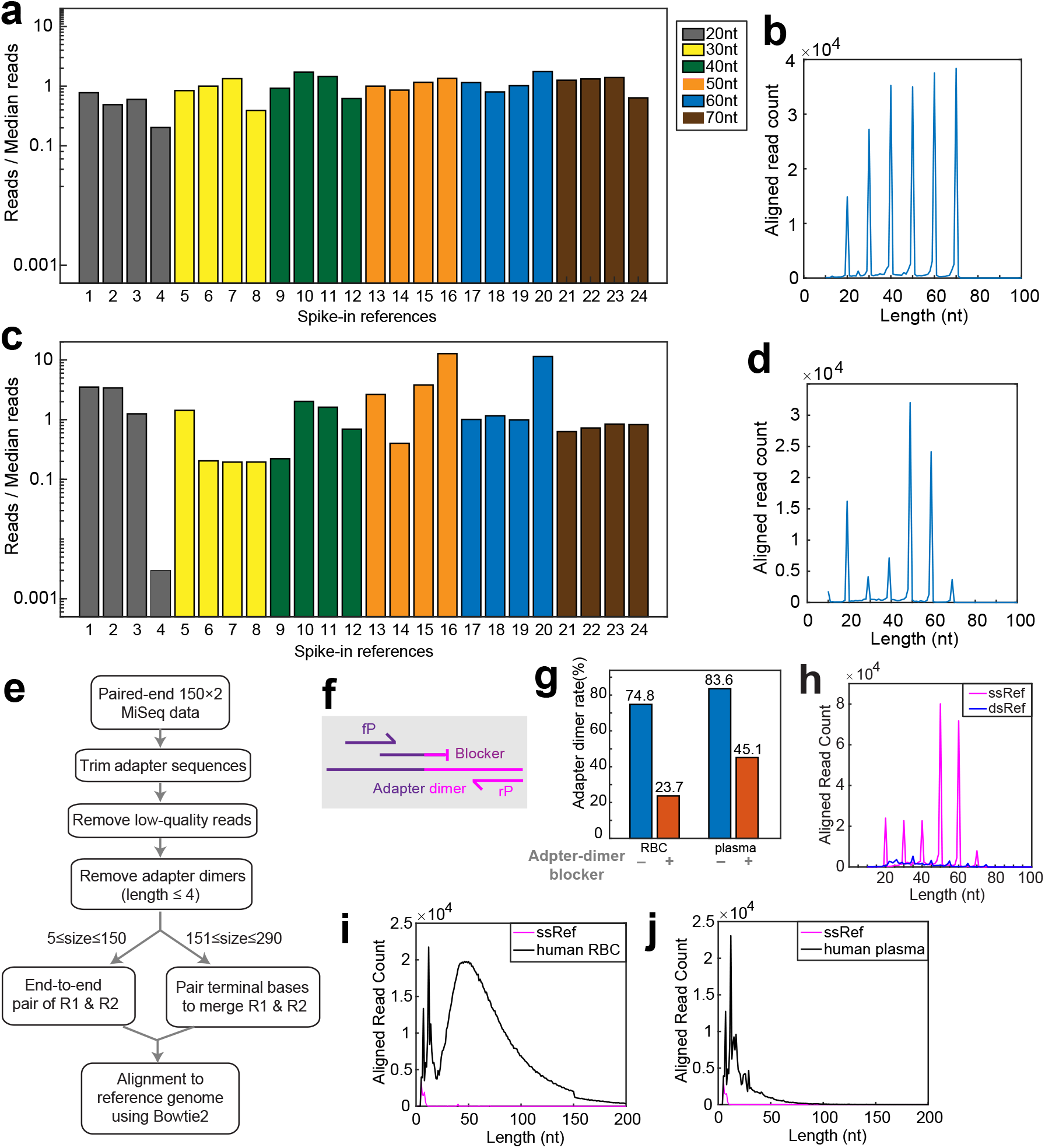
Workflow characterization. **(a)** Fold difference of reads aligned to each 1pM spike-in reference after library preparation using SRSLY PicoPlus kit. Reference strands are ssDNA of lengths from 20nt to 70nt with 10nt interval. 23/24 references are within 2-fold from the median reads. Length of reference ssDNA are color coded in the bar graph. **(b)** Length distribution of library prepared from spike-in ssDNA reference. 88.93% of aligned reads were recovered at expected sizes. **(c)** Fold difference of reads aligned to each 1pM spike-in reference after ussDNA capture and library preparation using SRSLY PicoPlus kit. All reference materials are recovered in the sequencing library, albeit with reduced reads uniformity, suggesting variations in capture yield. 22/24 references are within 10-fold from the median reads. **(d)** Length distribution of library prepared from captured spike-in ssDNA reference. Sharp peaks at expected sizes suggest unbiased length representation, and 86.26% of aligned reads are at expected sizes. **(e)** Bioinformatics pipeline for sequencing data analysis. **(f)** Design schema of adapter dimer blocker. Purple lines: i5 adapter sequences, pink lines: i7 adapter sequences, fP: forward primer, rP: reverse primer. **(g)** Adapter dimer rate of libraries prepared from RBC or plasma with (orange bars) and without (blue bars) adapter dimer blocker. **(h)** Reads aligned to ssDNA (magenta) and dsDNA (blue) references at equal molarity of 1pM/strand. dsDNA references are sized at 25:10:75 nt with 4 different strands at each size. 9x more reads are aligned to ssDNA reference than dsDNA reference, suggesting capture preference for ssDNA. **(i-j)** Length distribution of ussDNA in **(i)** RBC and **(j)** plasma from no-probe capture. Experiments were performed following standard direct capture and sequencing workflow except that LNA probes were not added.

**Figure S2:**
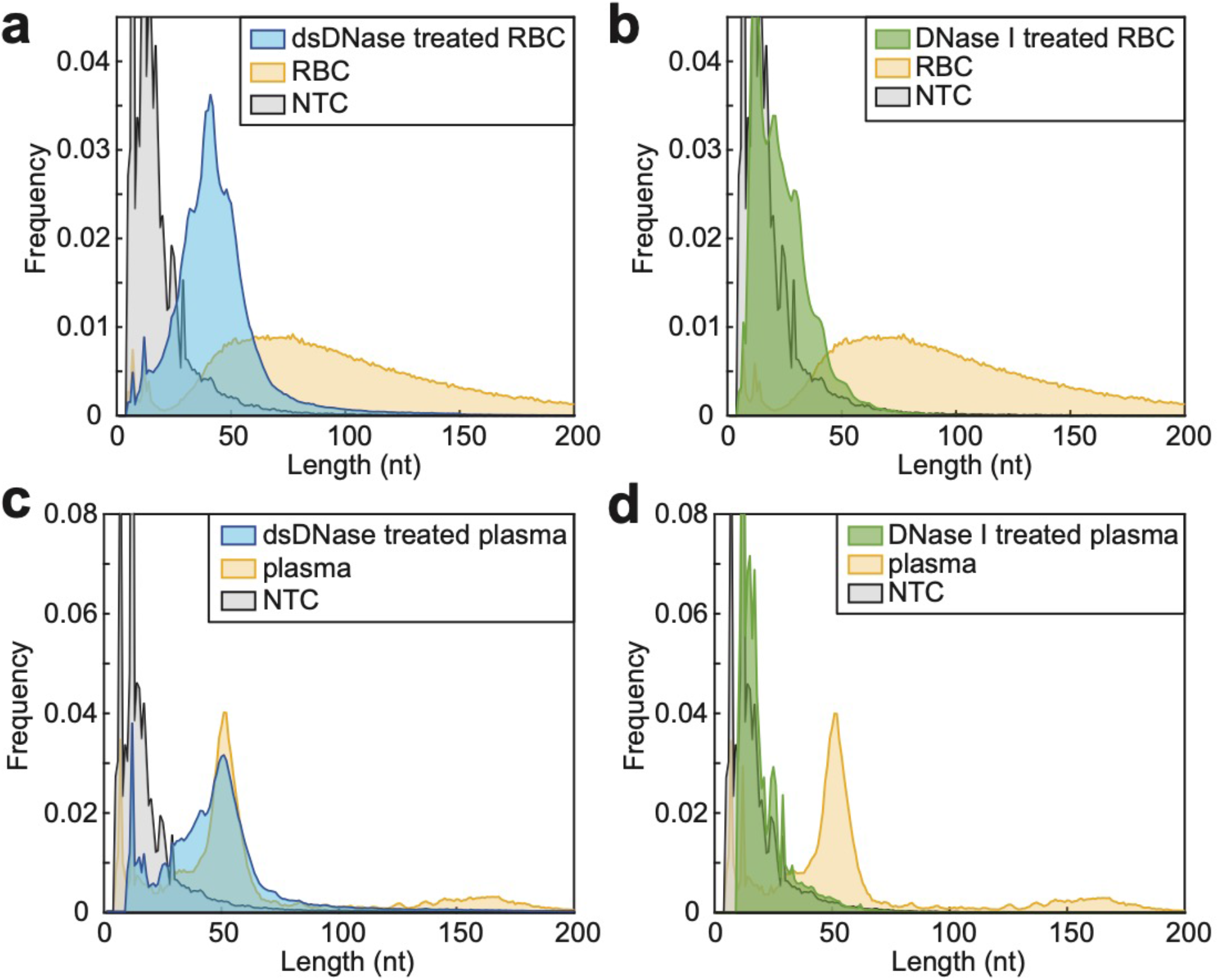
Blood-derived ussDNA underwent enzymatic digestion. Length distribution of **(a)** dsDNase treated RBC ussDNA, **(b)** DNase I treated RBC ussDNA, **(c)** dsDNase treated plasma ussDNA and **(d)** DNase I treated plasma ussDNA compared with untreated RBC or plasma and no template control (NTC). Yellow shades: untreated ussDNA in RBC or plasma; grey shades: NTC; Blue shades: ussDNA in RBC or plasma treated by dsDNase; Green shades: ussDNA in RBC or plasma treated by DNase I. dsDNase specifically digests double-stranded DNA and DNase I cleaves all DNA structures into short fragments.

**Figure S3:**
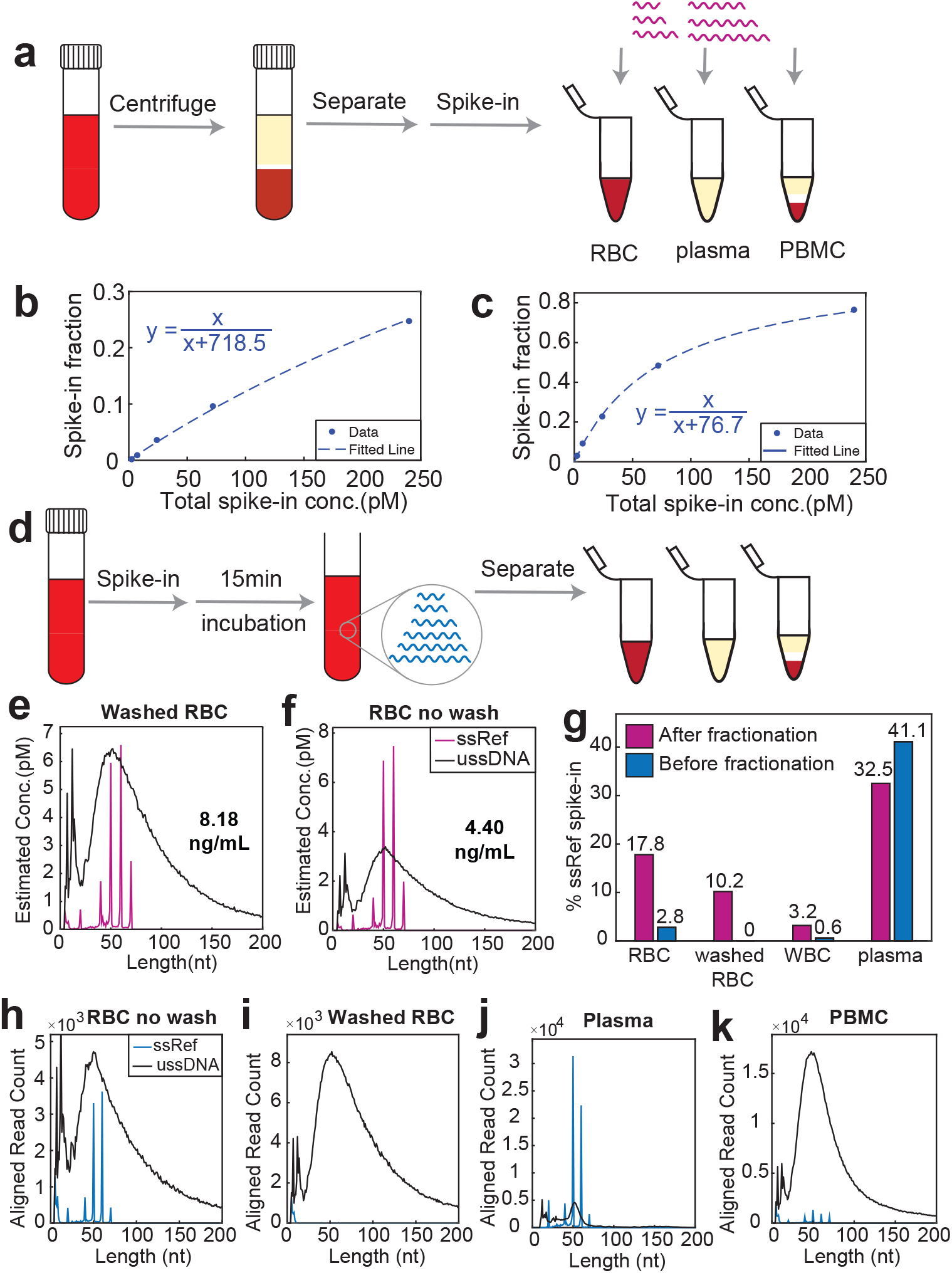
Spike-in of short synthetic DNA oligos as reference. **(a)** Schematic workflow of spiking in ssDNA strands to fractionated blood for quantitating blood-derived ussDNA. ssDNA references are composed of 24 strands at sizes 20:10:70 nt with 4 different strands at each size. **(b-c)** Calibration of spike-in fraction vs. total spike-in concentration to model ussDNA concentration in **(b)** RBC and **(c)** plasma. ssDNA references were spiked in at 0.1pM, 0.3pM, 1pM, 3pM and 10pM per strand and 24 strands in total. Dashed line stands for curve fitting of spike-in data points to y = x/(x+k) where y = spike-in fraction, x = total spike-in concentration (pM) and k is fitting coefficient representing ussDNA concentration in pM. The estimated ussDNA molarity is 718.5pM in RBC and 76.7pM in plasma. **(d)** Schematic workflow of spiking in ssDNA strands to total blood followed by fractionation. **(e-f)** Length distribution and concentration of ussDNA in **(e)** washed RBC (RBC pellet washed twice in PBS buffer) and **(f)** non-washed RBC. **(g)** Comparison of spike-in fraction between ssDNA strands added before (blue bars) and after (magenta bars) whole blood was fractionated. **(h-k)** Length distribution of ussDNA and ssDNA reference in blood fractionated after whole blood spike-in. Whole blood was fractionated into **(h)** RBC **(i)** washed RBC **(j)** plasma and **(k)** PBMC.

**Figure S4:**
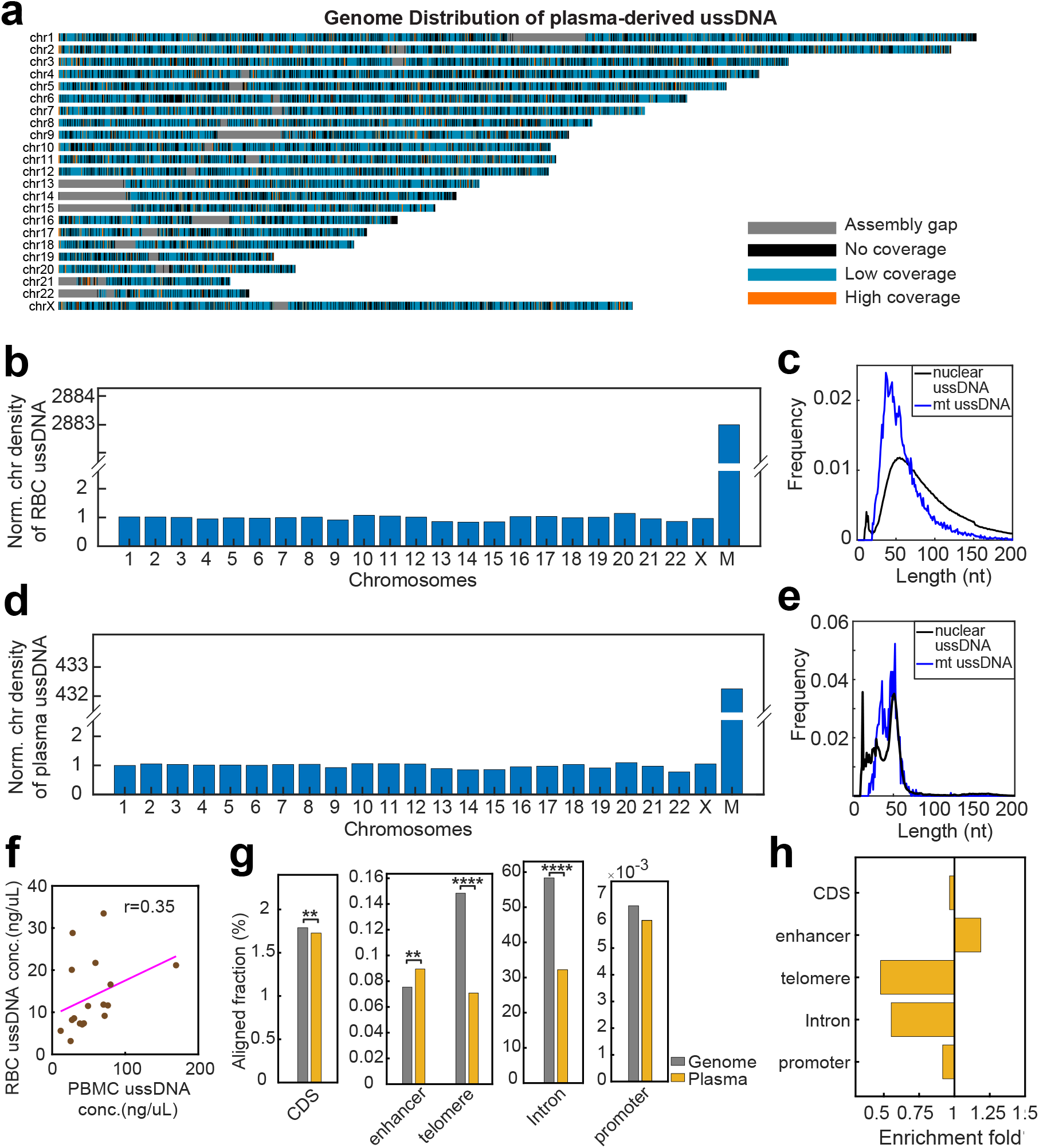
Genome distribution of blood-derived ussDNA. **(a)** Human genome distribution of plasma-derived ussDNA in autosomes and chromosome X. Regions where no reads aligned to were shown in black; gaps in human genome assembly GRCh38 were colored in grey. Regions with more than 2x global density were colored in orange. **(b)** RBC ussDNA density in chromosomes (including mitochondria genome, ChrM) normalized to global density. **(c)** Size distribution of mitochondrial ussDNA and nuclear ussDNA in RBC. **(d)** Plasma ussDNA density in chromosomes normalized to global density. 2880-fold and 430-fold enrichment of mitochondria genome are observed in RBC and plasma ussDNA, respectively. **(e)** Size distribution of mitochondrial ussDNA and nuclear ussDNA in plasma. **(f)** Correlation between RBC ussDNA concentration and PBMC ussDNA concentration. Pearson’s correlation coefficient (r) showed weak positive correlation. **(g)** Fraction of plasma ussDNA aligned to functional elements compared to their genome fractions. CDS: gene coding region. Plasma ussDNA exhibited significant enrichment of enhancer, and depletion in telomere, CDS and intron regions compared to random distribution (^**^ p<0.01, ^****^ p<0.0001). **(h)** Fold enrichment of functional elements in ussDNA derived from plasma.

**Figure S5:**
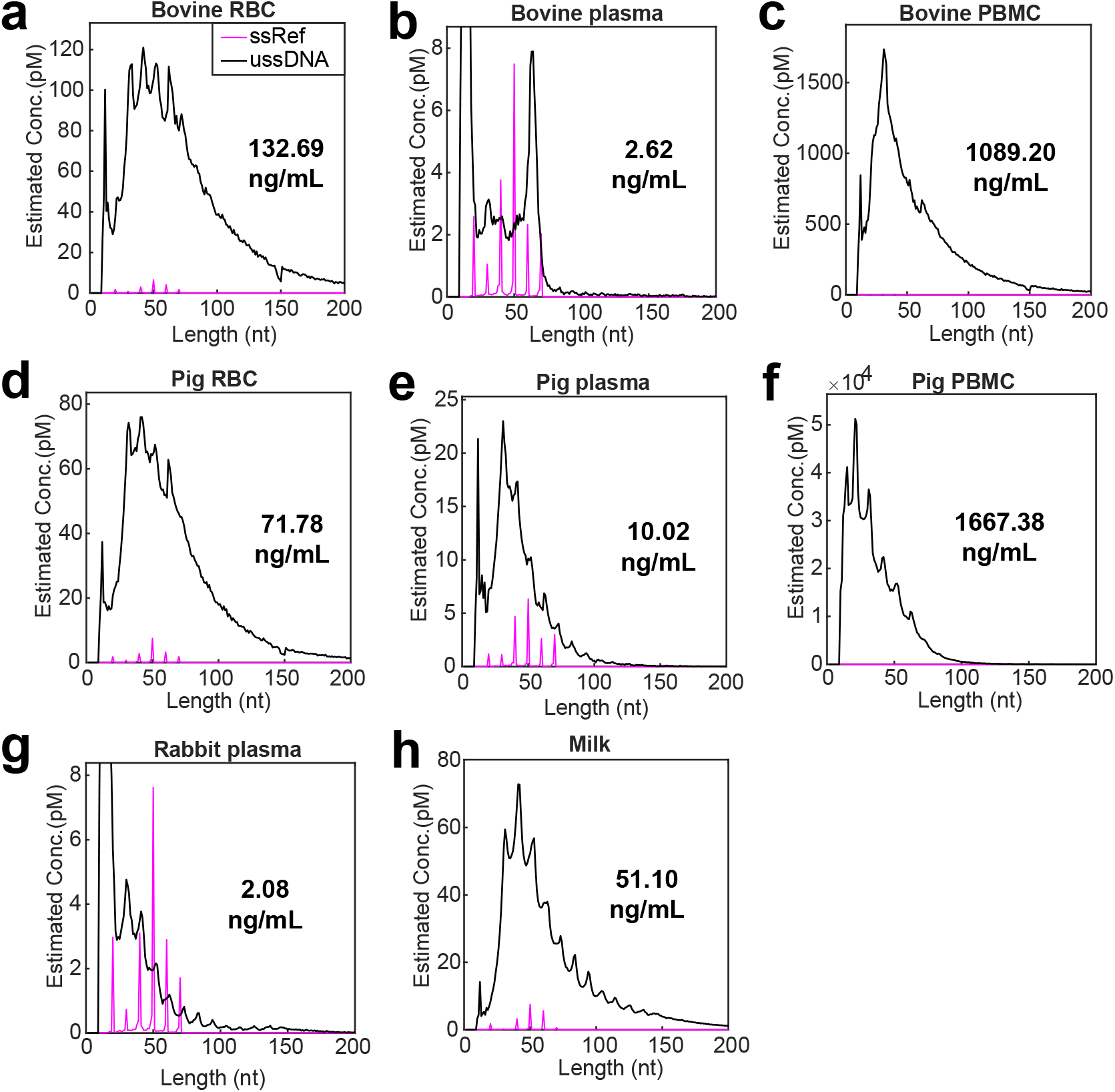
Size distributions and concentrations of ussDNA found in animal biofluids including **(a)** bovine RBC, **(b)** bovine plasma, **(c)** bovine PBMC, **(d)** pig RBC, **(e)** pig plasma, **(f)** pig PBMC, **(g)** rabbit plasma, and **(h)** milk. Magenta spikes represent ssDNA spike-in references, 4 different strands were spiked in at each size at concentration of 1pM each. Peaks below 20nt appear in most specimens and are considered artifacts produced by the workflow.

**Figure S6:**
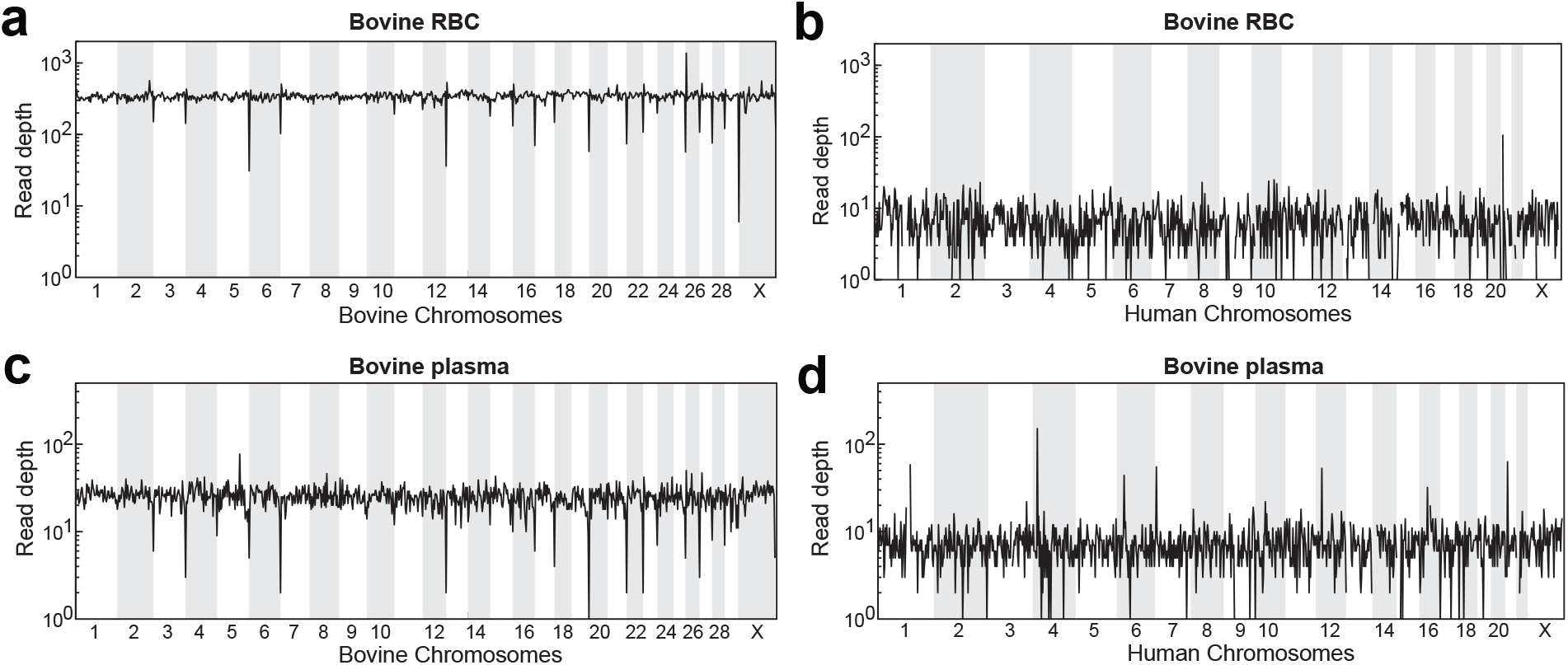
Cross-species genome alignment of bovine ussDNA. **(a)** Whole genome alignment of bovine RBC ussDNA to bovine genome. **(b)** Cross genome alignment of bovine RBC ussDNA to human genome. **(c)** Whole genome alignment of bovine plasma ussDNA to bovine genome. **(d)** Cross genome alignment of bovine plasma ussDNA to human genome. Each data point on the plotted line represents the number of reads aligned to 3 million nucleotide-sized bins. Cross-aligning bovine ussDNA to human genome showed notably reduced coverage.

**Figure S7:**
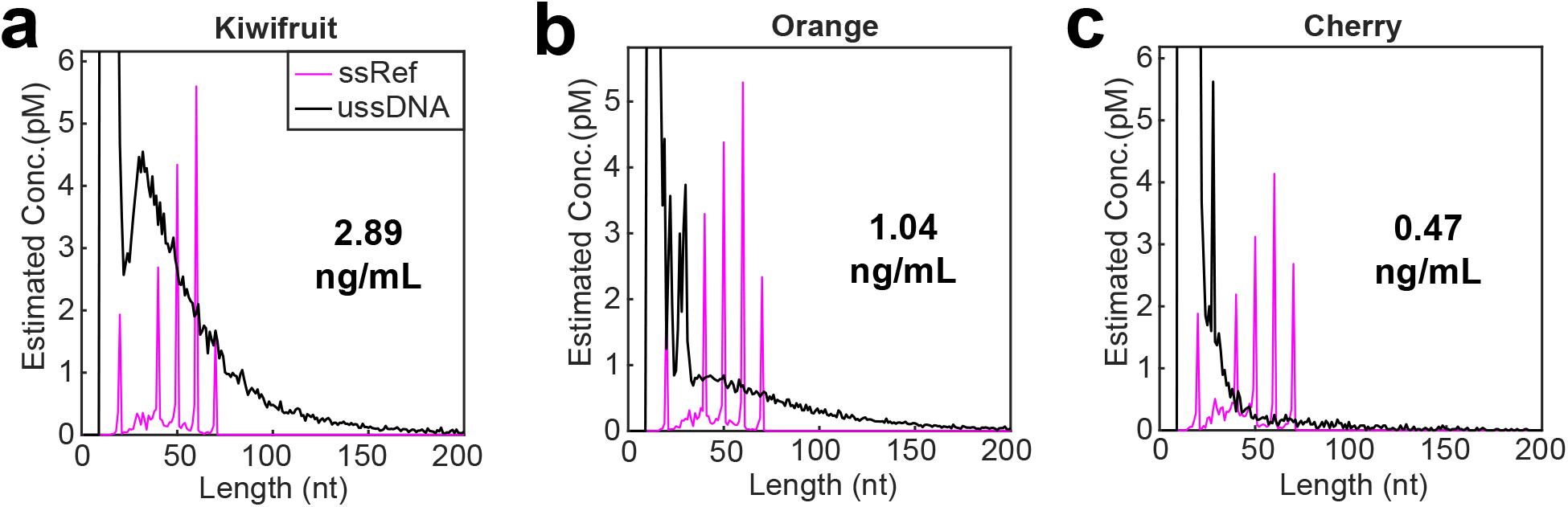
Size distributions and concentrations of ussDNA found in plant biofluids including **(a)** kiwifruit juice, **(b)** orange juice, and **(c)** cherry juice. Magenta spikes represent ssDNA spike-in references, 4 different strands were spiked in at each size at concentration of 1pM each.

## Methods

### Collection of biospecimen

Total blood samples were collected into K2EDTA collection tubes (BD #367863). N=7 of total blood samples were collected from healthy volunteers by a certified phlebotomist through venipuncture following IRB-FY2018-426 approved by Rice University. N=10 human blood samples were purchased from ZenBio, shipped on collection day at 4°C and delivered next day. Demographic information of blood donors was provided by ZenBio. Animal total blood samples were purchased from Discovery Life Sciences, shipped on collection day at 4°C and delivered next day. Demographic information of all 17 subjects was summarized in **Table. S1**.

**Table. S1.** Demographic information of healthy blood donors.

| No. | Collection type | Age | Gender |
| --- | --- | --- | --- |
| 1 | On-site | 26 | Female |
| 2 | On-site | 25 | Female |
| 3 | On-site | 25 | Male |
| 4 | On-site | 24 | Male |
| 5 | On-site | 25 | Male |
| 6 | On-site | 40 | Male |
| 7 | On-site | 34 | Female |
| 8 | ZenBio purchase | 60 | Male |
| 9 | ZenBio purchase | 26 | Female |
| 10 | ZenBio purchase | 38 | Male |
| 11 | ZenBio purchase | 44 | Female |
| 12 | ZenBio purchase | 31 | Female |
| 13 | ZenBio purchase | 47 | Female |
| 14 | ZenBio purchase | 28 | Male |
| 15 | ZenBio purchase | 69 | Male |
| 16 | ZenBio purchase | 49 | Male |
| 17 | ZenBio purchase | 35 | Male |

Milk was sampled from whole milk bought from grocery. Fruit juiced were collected via peeling and squeezing the fruits. Centrifugation at 1500xg for 10min was performed and only supernatant was collected.

### Blood fractionation

Within 2 hours following blood drawing from volunteers or upon receive of purchased blood samples, total blood samples were fractionated into plasma, peripheral blood mononuclear cell (PBMC) and red blood cell (RBC) fractions as follows. First, the total blood was centrifuged at 1800 xg for 15min with brake set to 1/3 of maximum level. Next, the upper clear layer of plasma was separated without disturbing the interface, and a p1000 pipettor was used to carefully collect PBMC layer to a new tube (*Residues of plasma and RBC remained in PBMC fraction because buffy coat cannot be cleanly separated). Then for the remaining content of condensed red blood cell pellet, 2x volume of PBS was gently added and tube was inverted ten times for mixing. The wash buffer was then removed following centrifuging at 500 xg for 5 minutes and discarding the supernatant. The washing step was repeated and a final centrifugation at 1500 xg for 5min was performed. The washed RBC was collected by discarding supernatant and transferring the pellet to a new tube. Unless otherwise mentioned, RBC fraction in this manuscript refers to RBC following 2x wash in PBS buffer.

### Direct capture

Immediately following blood fractionation, 100µL of fractionated blood components were mixed with LNA capture probe (5’-+N+N+N+N+N+N+N+N+N+N/Sp18/Bio/-3’, Integrated DNA Technologies) and hybrid capture buffer at final concentrations of 2µM capture probe, 0.5M NaCl, 1x TE and 0.1% tween20. The capture mixture was briefly vortexed and incubated at room temperature for 2 hours with gentle shaking. For each sample, 60µL of Streptavidin beads (Thermal Fisher, #65001) that pre-equilibrated to room temperature were pelleted using a magnetic rack and resuspended in 10x volume of 0.5M NaCl solution to wash the beads. Beads were re-pelleted on the magnetic rack to remove the wash buffer and resuspended in 100µL buffer of 0.5M NaCl, 1x TE and 0.1% tween20. The buffer was removed by separating streptavidin beads on a magnetic rack, and the pelleted beads were suspended in hybrid capture mixture and incubated at room temperature for 30min to allow binding of biotinylated probes to beads. The streptavidin beads were then washed three times in 500µL of 0.5M NaCl, 1x TE and 0.1% tween20. To collect bound DNA, beads were resuspended in 25µL of 0.1x TE buffer and heated at 95°C for 5min to dissociate captured DNA from LNA probe. Eluant containing captured DNA was transferred to a PCR tube following pelleting beads on magnetic rack.

### ussDNA library preparation

Library preparation of captured single-stranded DNA was performed using SRSLY NanoPlus (Claret Biosciences, #CBS-K250B-96) according to manufacturer’s instructions with modifications to reduce adapter dimer. Specifically, 2µL of ss Enhancer and 18µL of eluted ssDNA were mixed and heated at 98°C for 3min and immediately cooled to 4°C. Then 2µL of SRSLY NGS Adapter A, 2µL of SRSLY NGS Adapter B and 26µL of SRSLY Master Mix were added and the mixture was incubated in a thermocycler at 37°C for 1hr with lid temperature set to 45°C. The product was purified with AMPure XP beads (Beckman Coulter, #A63881) and isopropanol was added to increase recover of short fragments at ratio of [reaction product : AMPure beads : water : isopropanol = 50µL : 59.4µL : 48.4µL : 11.6µL]. Then the library was PCR-amplified with custom designed PCR oligos (forward primer: 5’-ACACTCTTTCCCTACACGACG-3’; reverse primer: 5’-GTGACTGGAGTTCAGACGTGT-3’) and adapter dimer blocker (5’-CACGACGCTCTTCCGATCTAGATCG/3SpC3/-3’) that selectively suppress amplification of adapter dimers. The PCR was performed with 400nM of each primer and 4µM of adapter dimer blocker in PowerUp SYBR Green Master Mix (Thermal Fisher, #A25742), and with thermocycle program of initiation at 95°C for 3min and 13 cycles of 95°C for 10s and 60°C for 40s (short as 95°C 3min – (95°C 10s - 60°C 40s) x13). After purification with Monarch PCR & DNA Cleanup kit (NEB, #T1030S), libraries were indexed using NEBNext Multiplex Oligos (NEB, #E7780S) and Taq Universal probes supermix (Bio-Rad, #1725131) for 8 cycles of PCR following 95°C 3min – (95°C 10s - 60°C 40s) x8. The indexed library was purified with 1.5x volume of AMPure beads and quality controlled by bioanalyzer. Libraries were sequenced on an illumina Miseq or Nextseq instrument using 150×2 chemistry. 1-3 million reads were allocated to each sample, representing 1/2000 coverage of human genome.

### Enzymatic digestion of captured ussDNA

ussDNA extracted from RBC or plasma were treated with dsDNase (Thermal Fisher, #EN0771) to specifically digest double-stranded DNA and DNase I (NEB, #M0303S) to digest both double-stranded DNA as well as single-stranded DNA. Specifically, after binding captured ussDNA to streptavidin-coated magnetic beads, the beads were washed to remove biofluids and tissue debris. Washed streptavidin beads were pelleted using a magnetic beads and wash buffer discarded. 2μL of enzyme, 5μL of 10x buffer and 43μL of water were mixed and used to suspend DNA-bound streptavidin beads. The mixture was incubated at 37°C for 5min for dsDNase treatment or at 37°C for 10min for DNase treatment incubate. Following enzymatic digestion, beads were pelleted again and washed once in 200μL of 10 mM Tris-HCl, 1 mM EDTA, 0.05% Tween-20, 100 mM NaCl, 0.5% SDS, and another time in 200μL of 10 mM Tris-HCl, 1 mM EDTA, 0.05% Tween-20, 100 mM NaCl. DNA molecules were then released from beads by heat incubation at 95°C for 5min in 0.1x TE buffer.

### Reference material preparation

Reference materials were synthetic short single-stranded oligos purchased from Integrated DNA Technologies (IDT). Sequences of reference materials are generated by a custom random generator with GC% range between 40% and 60% and sequence complexity resemble biological sequences. Homology test was implemented to ensure that the sequences did not align to human genome. Single-stranded spike-in references are composed of 24 different oligos at equal concentrations with sizes of 20nt, 30nt, 40nt, 50nt, 60nt and 70nt, with four distinct sequences at each size. Double-stranded spike-in references are composed of 24 oligo duplexes at equal concentrations with sizes of 25nt, 35nt, 45nt, 55nt, 65nt and 75nt, with four distinct sequences at each size. To form oligo duplexes, each two complementary oligos were annealed at 10μM in 1x PBS buffer to form a single double-stranded species using a thermal annealing program of denaturing at 95°C for 5min followed by cooling to 20°C at rate of -0.1°C/6s, and stored at -20°C until use.

### ussDNA concentration quantitation

24 ssDNA oligos were spiked into hybrid capture mixture (Hyb) at final concentration of 1pM/strand to be used as a reference to estimate concentration of blood-derived ussDNA. Let R_genome_ denote number of reads aligned to reference genome with size >20nt, and R_spikein_ denote number of reads aligned to all single-stranded spike-in sequences. The formula for estimating the overall concentration of ussDNA is the following:

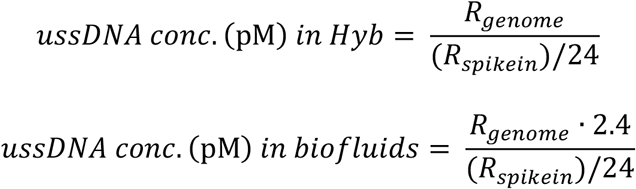

*Multiplied by 2.4 because when 100μL of biofluids was added to hybridization mixture, its concentration was diluted by 2.4-fold.

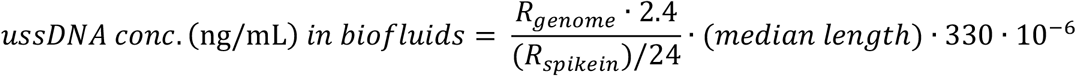

### Bioinformatics analysis

Fastq files were pre-processed using custom python scripts as follows. Fastq sequences are trimmed off of adapter sequences, and then quality filtered by retaining reads with greater than 80% of high quality (Q30) bases. Reads with sizes less than 4 bases are considered adapter dimers and was removed from further analysis. Short sequences (<150nt) are end-to-end paired and bases with higher quality between read1 and read2 are retained. Longer reads (150nt<length<290nt) cannot be length-resolved by a single read and thus match of terminal sequences of read1 and read2 is performed to merged matched reads. Next, filtered fastq reads are aligned to reference genome using Bowtie2 to generate sam files containing alignment information (**Fig. S1e**). Then custom MATLAB scripts were used to analyze sequence size and distribution.

### Genome distribution analysis

Genome distribution analysis is based on genomic alignment coordinates given by sam files. For global genome distribution analysis, the GRCh38 genome assembly was separated into bins of 10,000 bp, and the number of reads aligned to each bin was calculated and the coverage was plotted as a heatmap cross all chromosomes (excluding Y chromosome). Chromosome bias was analyzed by counting reads aligned to each chromosome including the mitochondrial genome (chrM). The chromosomal density was then calculated by dividing aligned reads by chromosome size, which was then normalized to global density. Genomic locations of functional elements including gene coding regions, introns, telomeres, promoters and enhancers were downloaded from UCSC genome browser. Here, telomeres are defined as most distal 1M bases flanking all chromosomes. Reads with overlapping coordinates were considered the corresponding functional element. Statistical analysis of the distribution of functional elements assumes that the reads aligned to each element is proportional to the element size. Each read was considered a trial in binomial distribution with the probability being the fraction of genome occupied by certain element. Therefore, for each functional element, significance level could be calculated from binomial distribution with probability and aligned reads.

## Data availability

The data supporting the findings of this study are available within the paper. Source data are available upon request.

## Code availability

The custom Python and MATLAB scripts used in this study are available as Supplementary Software.

## Acknowledgements

This work was supported by NIH grant no. R01CA233364 to D.Y.Z. We thank Dr. Ping Song for supervising phlebotomy practice.

## References

1. Snyder, M.W., Kircher, M., Hill, A.J., Daza, R.M. & Shendure, J. Cell-free DNA comprises an in vivo nucleosome footprint that informs its tissues-of-origin. Cell 164, 57–68 (2016).

2. Lo, Y.D. et al. Maternal plasma DNA sequencing reveals the genome-wide genetic and mutational profile of the fetus. Science translational medicine 2, 61ra91–61ra91 (2010).

3. Wyatt, A.W. et al. Concordance of circulating tumor DNA and matched metastatic tissue biopsy in prostate cancer. JNCI: Journal of the National Cancer Institute 109 (2017).

4. Chae, Y.K. et al. Concordance between genomic alterations assessed by next-generation sequencing in tumor tissue or circulating cell-free DNA. Oncotarget 7, 65364 (2016).

5. Szpechcinski, A. et al. Cell-free DNA levels in plasma of patients with non-small-cell lung cancer and inflammatory lung disease. British journal of cancer 113, 476–483 (2015).

6. Meddeb, R. et al. Quantifying circulating cell-free DNA in humans. Scientific reports 9, 1–16 (2019).

7. Mouliere, F. et al. Enhanced detection of circulating tumor DNA by fragment size analysis. Science translational medicine 10 (2018).

8. Liu, X. et al. Enrichment of short mutant cell-free DNA fragments enhanced detection of pancreatic cancer. EBioMedicine 41, 345–356 (2019).

9. Streubel, A. et al. Comparison of different semi-automated cfDNA extraction methods in combination with UMI-based targeted sequencing. Oncotarget 10, 5690 (2019).

10. Cook, L. et al. Does size matter? Comparison of extraction yields for different-sized DNA fragments by seven different routine and four new circulating cell-free extraction methods. Journal of clinical microbiology 56, e01061–01018 (2018).

11. Kierzek, E. et al. The influence of locked nucleic acid residues on the thermodynamic properties of 2′-O-methyl RNA/RNA heteroduplexes. Nucleic acids research 33, 5082–5093 (2005).

12. Troll, C.J. et al. A ligation-based single-stranded library preparation method to analyze cell-free DNA and synthetic oligos. BMC genomics 20, 1–14 (2019).

13. Zhang, J.X. et al. Predicting DNA hybridization kinetics from sequence. Nature chemistry 10, 91–98 (2018).

14. Wu, L.R., Chen, S.X., Wu, Y., Patel, A.A. & Zhang, D.Y. Multiplexed enrichment of rare DNA variants via sequence-selective and temperature-robust amplification. Nature biomedical engineering 1, 714–723 (2017).

15. Jiang, P. et al. Lengthening and shortening of plasma DNA in hepatocellular carcinoma patients. Proceedings of the National Academy of Sciences 112, E1317–E1325 (2015).

16. Kelly, R.D., Mahmud, A., McKenzie, M., Trounce, I.A. & St John, J.C. Mitochondrial DNA copy number is regulated in a tissue specific manner by DNA methylation of the nuclear-encoded DNA polymerase gamma A. Nucleic acids research 40, 10124–10138 (2012).

